# Divergent selection and drift shape the genomes of two avian sister species spanning a saline-freshwater ecotone

**DOI:** 10.1101/344614

**Authors:** Jennifer Walsh, Gemma V. Clucas, Matthew D. MacManes, W. Kelley Thomas, Adrienne I. Kovach

**Affiliations:** Department of Natural Resources and the Environment, University of New Hampshire, Durham, NH 03824, USA; Fuller Evolutionary Biology Program, Cornell Laboratory of Ornithology, Cornell University, Ithaca, NY 14850, USA; Department of Ecology and Evolutionary Biology, Cornell University, Ithaca, NY 14853, USA; Department of Molecular, Cellular and Biomedical Sciences, University of New Hampshire, Durham, NH 03824, USA; Hubbard Center for Genome Studies, Durham, NH 03824, USA

**Keywords:** *Ammodramus caudacutus*, *Ammodramus nelsoni*, Genomics, Ecological Divergence, Ecological Speciation, Demography, Tidal marshes, Adaptation

## Abstract

The role of species divergence due to ecologically-based divergent selection – or ecological speciation – in generating and maintaining biodiversity is a central question in evolutionary biology. Comparison of the genomes of phylogenetically related taxa spanning a selective habitat gradient enables discovery of divergent signatures of selection and thereby provides valuable insight into the role of divergent ecological selection in speciation. Tidal marsh ecosystems provide tractable opportunities for studying organisms’ adaptations to selective pressures that underlie ecological divergence. Sharp environmental gradients across the saline-freshwater ecotone within tidal marshes present extreme adaptive challenges to terrestrial vertebrates. Here we sequence 20 whole genomes of two avian sister species endemic to tidal marshes – the Saltmarsh Sparrow (*Ammodramus caudacutus*) and Nelson’s Sparrow (*A. nelsoni*) – to evaluate the influence of selective and demographic processes in shaping genome-wide patterns of divergence. Genome-wide divergence between these two recently diverged sister species was notably high (genome-wide F*_ST_* = 0.32). Against a background of high genome-wide divergence, regions of elevated divergence were widespread throughout the genome, as opposed to focused within islands of differentiation. These patterns may be the result of genetic drift acting during past tidal march colonization events in addition to divergent selection to different environments. We identified several candidate genes that exhibited elevated divergence between Saltmarsh and Nelson’s sparrows, including genes linked to osmotic regulation, circadian rhythm, and plumage melanism – all putative candidates linked to adaptation to tidal marsh environments. These findings provide new insights into the roles of divergent selection and genetic drift in generating and maintaining biodiversity.

## Introduction

The role of differential adaptation to divergent selective environments in generating and maintaining biodiversity has become an increasing focus for evolutionary biologists over the past decade and has been termed ecological speciation (Schluter 2001; Rundle and Nosil, 2005; Funk et al. 2006). While the concept of ecologically-based divergent selection is not new (Mayr 1942; Rundle and Nosil 2005), disentangling the contribution of ecological forces from non-ecologically based evolutionary forces (i.e., reductions in population size, genetic drift) remains a challenge. Despite these challenges, increasing accessibility and improvement of current sequencing technologies have allowed for application of whole-genome sequencing to questions in natural populations (Ellegren 2014; Toews et al. 2016; Campagna et al. 2017). The genomics era holds promise for ecological speciation research, as it has allowed for the detection of divergent signatures of selection on a genome-wide scale. In recent years, the application of genomics to non-model systems has provided new insight into mechanisms underlying lineage-specific adaptations in a range of taxa (Qiu et al. 2012; Cai et al. 2013; Zhan et al. 2013, Liu et al. 2015; Li et al. 2017). It has also provided insight into the genomic architecture of adaptation and speciation (Strasburg et al. 2011, Cruickshank and Hahn, 2014, Larson et al. 2017).

Under a scenario of ecological speciation, divergent selection will result in differential performance of individuals inhabiting alternative ecological niches (Nosil 2012; Arnegard et al. 2014). Reproductive isolation resulting from such adaptive divergence may occur either in sympatry, parapatry, or allopatry (Nosil, 2007, 2012; Langerhans et al. 2007). Comparison of the genomes of two ecologically divergent taxa spanning a selective habitat gradient provides valuable insight into the role of niche divergence in driving natural selection and speciation within a system. Characterizing genomic differentiation in recently diverged taxa is critical for increasing understanding of ecological speciation, as early-acting barriers to gene flow have a larger effect on driving reproductive isolation than late-acting barriers (Coyne and Orr 2004). In addition, identifying elevated regions of differentiation – and potential genomic regions under selection – is easier when baseline divergence is low, as expected for phylogenetically closely related and recently diverged taxa (Ellegren et al. 2012; Poelstra et al. 2014; Toews et al. 2016; Campagna et al. 2017).

Tidal marsh habitats in North America have undergone rapid changes since the last glacial maximum (Greenberg et al. 2006), and they are comprised of sharp ecotones that provide highly tractable opportunities for understanding underlying spatial patterns of genetic variation and adaptation. Because most tidal marsh endemics have colonized these habitats only after the rapid expansion of coastal marshes approximately 5,000 – 7,000 years ago (Malamud-Roam et al. 2006), these systems provide opportunities to investigate patterns of recent and contemporary evolution. Environmental gradients across the saline-freshwater ecotone present extreme adaptive challenges to terrestrial vertebrates (Greenberg 2006; Bayard et al. 2011). Divergent selection among populations spanning these salinity gradients can be apparent in both physiological (*i.e.*, pathways involved in osmotic regulation) and morphological (*i.e.*, plumage – saltmarsh melanism, bill and body size variation; Grinnell 1913; Grenier and Greenberg 2005) traits. These strong ecological gradients in tidal marshes provide a model system for applying comparative genomic analysis to investigate the role of ecological divergence in shaping species diversity.

Here we investigated patterns of genome-wide differentiation between two recently diverged marsh endemics, the Saltmarsh (*Ammodramus caudacutus*) and Nelson’s (*A. nelsoni*) sparrow (~600,000 years; Rising and Avise 1993). Although long considered a single species (AOU 1931), the Nelson’s and saltmarsh sparrow are currently recognized as two species comprised of a total of five subspecies: *A. nelsoni nelsoni* – breeds in the continental interior from eastern British Columbia to central Manitoba and northern South Dakota; *A. nelsoni alterus* – around the James and Hudson bays; *A. nelsoni subvirgatus* – across the Canadian Maritimes to southern Maine; *A. caudacutus caudacutus* – from southern Maine to New Jersey; and *A. caudacutus diversus* – from southern New Jersey to Virginia (Greenlaw and Rising 1994). The prevailing evolutionary hypothesis (Greenlaw 1993) suggests a history of vicariance for the saltmarsh and Nelson’s sparrow, whereby an ancestral population spanning a coastal to interior range was split by Pleistocene glaciation, resulting in an isolated interior population. Following differentiation, this interior population then spread eastward back toward the Atlantic coast after recession of the Wisconsin ice mass, making secondary contact with ancestral coastal populations, and establishing the current ranges and ecotypes within this species complex (Greenlaw 1993). Recent analyses of genetic and morphological characters indicate the strongest differences appear to correspond to habitat-type, clustering the five subspecies into three groups: 1) the two freshwater, interior subspecies of Nelson’s sparrow, 2) the brackish, coastal subspecies of Nelson’s sparrow, and 3) the two saltwater, coastal subspecies of saltmarsh sparrow (Walsh et al. 2017).

While both species inhabit tidal marshes in sympatry, variation in habitat affinity, morphology, and behavior suggest a role for divergent selection and adaptation in this system. Specifically, the saltmarsh sparrow is a narrow niche specialist that has been associated with salt marshes over a longer evolutionary time frame (possibly 600,000 years, Chan et al. 2006) compared to the Nelson’s sparrow, which uses a broader range of habitats including brackish and fresh water marshes and hay fields. Due to these differences in niche specificity, differentiation of these sister species may have been largely driven by divergent natural selection. Alternatively, changes in population size during vicariant isolation and colonization events may have increased the role of genetic drift in driving interspecific divergence. While previous work in this system has identified patterns consistent with divergent selection across the saline-freshwater ecotone (Walsh et al. 2015, 2016, 2017), the genome-wide pattern of differentiation between saltmarsh and Nelson’s sparrows remains unknown. An understanding of the genomic landscape of these taxa will reveal the influence of demographic processes and divergent selection in ecological speciation.

We sequenced whole genomes of saltmarsh and Nelson’s sparrows to investigate the role of divergent selection across an ecological gradient in shaping genome-wide patterns of divergence. We were interested in identifying genomic regions exhibiting elevated divergence due to selection across the saline – freshwater ecotone. We predicted elevated divergence between saltmarsh and Nelson’s sparrows in gene regions linked to known tidal marsh adaptations. Specifically, we hypothesized that genes linked to kidney development, osmotic regulation (salt tolerance in tidal environments; Greenberg et al. 2006; Goldstein 2006), circadian rhythm (important for nest initiation relative to tidal cycles; Shriver et al. 2007; Walsh et al. 2017), bill size (larger bills facilitate evaporative heat loss; Greenberg et al. 2012; Greenberg and Danner 2012; Luttrell et al. 2014), and melanic plumage (example of salt marsh melanism; Grinnell 1913; Greenberg and Droege 1990; Walsh et al. 2016) would be targets of selection in tidal marsh environments and would be key mechanisms underlying ecological divergence between these species.

## Methods

### Sample collection

For the reference genome, we sampled a male saltmarsh sparrow from the Marine Nature Center in Oceanside, New York in July 2016. Blood was collected from the brachial vein and stored in Puregene lysis buffer (Gentra Systems, Minneapolis, MN). For genome re-sequencing, we sampled 20 individuals, 10 Nelson’s sparrows (2 females and 8 males) and 10 saltmarsh sparrows (8 females and 2 individuals of unknown sex), from marshes along the northeastern coastline of the United States (Figure 1; Supplementary Information; table S1) during the breeding seasons (June – August) of 2008 – 2014. Nelson’s sparrows were sampled from three populations in Maine. Saltmarsh sparrows were sampled from ten populations in Massachusetts, Rhode Island, Connecticut, New York, Maine, and New Hampshire. From each bird, we collected blood samples from the brachial vein and stored them on Whatman filter cards.

**Figure 1:**
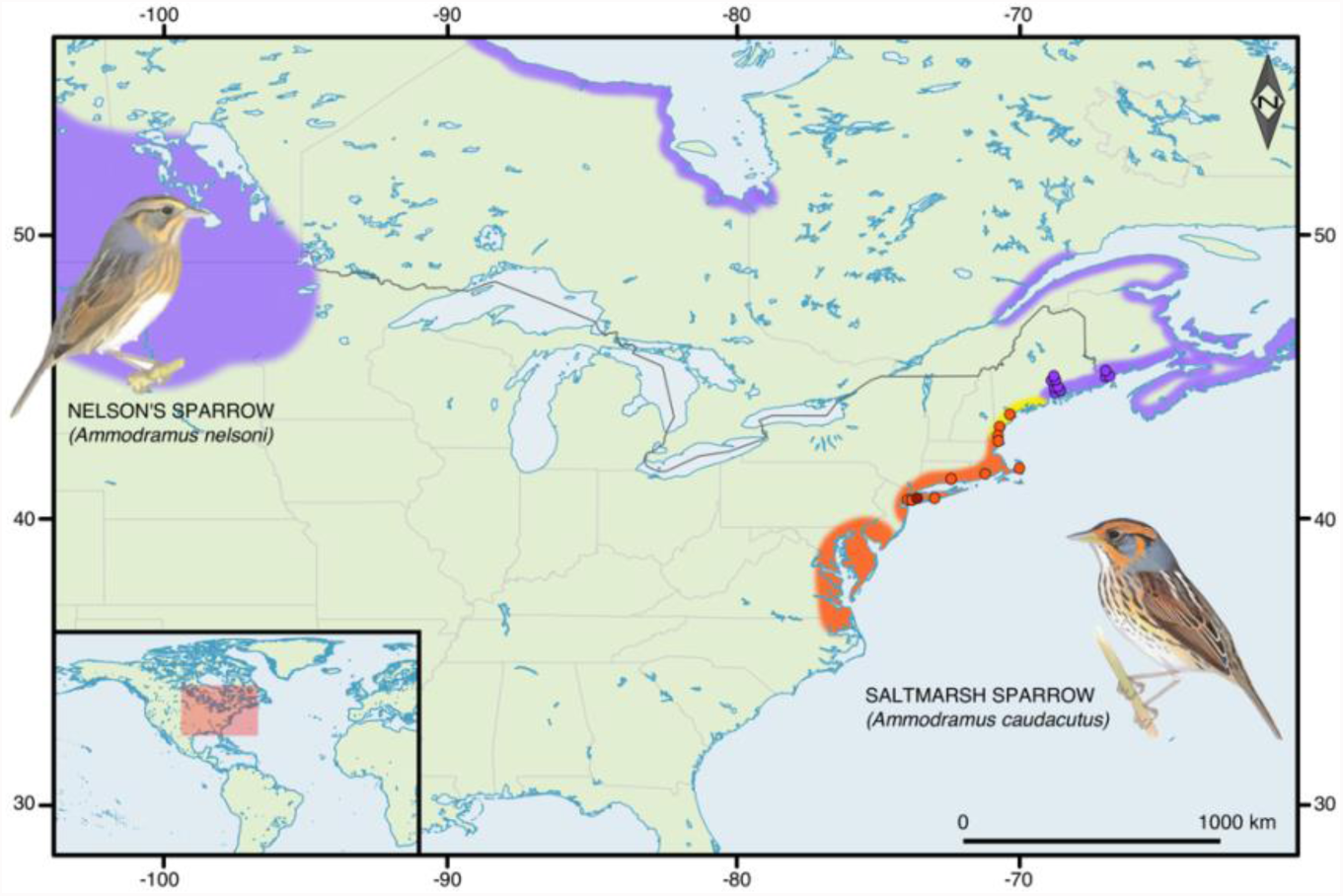
Breeding distributions and sampling locations. The breeding distributions of the Nelson’s and Saltmarsh sparrows are shown in purple and orange, respectively. The hybrid zone along the New England coastline is shown in yellow. The breeding locations of the Nelson’s sparrows that were resequenced in this study are shown by the purple dots, the resequenced Saltmarsh sparrows are shown by the orange dots, and the breeding location of the reference genome individual is shown in dark brown. The inset figures show the plumage of the two species.

### Reference Genome – Library Construction & Assembly

For reference genome sequencing, DNA was extracted from blood stored in lysis buffer using a PureGene extraction kit (Gentra, Minneapolis, MN). Three libraries were constructed using an Illumina Nextera library preparation kit: one paired-end (PE) library with a 180 bp insert size and two mate-pair (MP) libraries with 3 kbp and 8 kbp insert sizes. Each library was sequenced on a single HiSeq 2500 lane PE x 100 cycles by the Weill Cornell Medical College Core Genomics facility.

Genome assembly was performed with ALLPATHS-LG version-44849 (Gnerre et al. 2011; Ribeiro et al. 2012) using the default parameters. ALLPATHS-LG takes raw data as input, without prior adapter removal and trimming. We used Bioanalyzer results to estimate the insert-size and expected standard deviation, which are required input for ALLPATHS-LG. The assembly was completed in six days on a 64-core computer (1024GB RAM, 19TB hard drive) from the Cornell Computational Biology Service Unit BioHPC Lab. We obtained assembly statistics with Quast version 2.3.

### Reference Genome – Annotation

We annotated the saltmarsh sparrow assembly with the MAKER pipeline v 2.31.9 (Cantarel et al. 2008). Gene models were created using the zebra finch (*Taeniopygia guttata*) Ensemble protein database (downloaded March 2^nd^, 2017 from http://useast.ensembl.org/Taeniopygia_guttata/Info/Index?redirect=no) and a saltmarsh sparrow transcriptome. To generate the transcriptome, cDNA libraries were prepared from RNA extracted with a RNeasy kit (Qiagen, Valencia, CA, USA) from six tissues after overnight freezing – heart, muscle, liver, brain, gonad, and kidney – from a single male Saltmarsh Sparrow individual. Sequencing libraries were generated from polyA enriched mRNA using the Illumina TruSeq RNA sample prep LT system. RNA sequencing of 100 bp PE reads was performed in a single Illumina HiSeq2000 lane at Vanderbilt University. Transcriptome assembly was performed on the combined datasets with CLC Genomics Workbench (V5.1.2.) using CLC assembly Cell 4.0 set to default parameters. Genes were predicted with SNAP v2013-11-29, using an iterative training process inside MAKER v2.31.9, and Augustus v3.2.2_4, using a Hidden Markov Model from the chicken. This produced a total of 15,414 gene models, which included 85.2% complete BUSCO v2.0 (Simão et al. 2015) genes and a further 9.3% which were fragmented, when assessed against the aves_odb9 lineage data and specifying the white-throated sparrow (*Zonotrichia albicollis*) as the most closely related species in the database. The saltmarsh sparrow assembly was aligned to the zebra finch genome using *Satsuma* Chromosembler (Grabherr et al. 2010). Using this approach, scaffolds were mapped to chromosome coordinates via synteny. Thus, scaffolds were assigned to chromosomes for removal of sex-linked markers, however, population genetic analyses were conducted at the scaffold level.

### Whole-Genome Resequencing and Variant Discovery

We sequenced the genomes of an additional 20 individuals, 10 saltmarsh sparrows and 10 Nelson’s sparrows. DNA was extracted using the DNeasy blood and tissue kit (Qiagen, Valencia, CA, USA). For 7 of the 20 individuals (six saltmarsh and one Nelson’s Sparrow), we prepared individually barcoded Illumina TruSeq DNA libraries, from 1.2 to 31 µg of DNA, which were sequenced on a Illumina HiSeq 2000 at Vanderbilt University. DNA from one Saltmarsh Sparrow (31 µg) and one Nelson’s Sparrow (6.2 µg) individuals were sequenced each in a single lane, and five Saltmarsh Sparrows (1.2-3 µg DNA) were sequenced together in a third lane. For the remaining 13 individuals, we prepared individually barcoded libraries using 1-2 ng of DNA following the Nextera^®^ DNA library preparation kit protocol, with a target insert size of 550 bp. We pooled the 13 libraries using concentrations of adapter-ligated DNA determined through digital PCR and sequenced them as 150-bp PE reads in two lanes on an Illumina HiSeq2500 at the University of New Hampshire Hubbard Center for Genome Studies. The quality of individual libraries was assessed using FastQC version 0.11.5 (http://www.bioinformatics.babraham.ac.uk/projects/fastqc).

We used a combination of programs to perform sequence trimming, adapter removal, and quality filtering. These included seqtk v 1.1-r91 (https://github.com/lh3/seqtk/blob/master/README.md) and Skewer v 0.1.127 (Jiang et al. 2014). We allowed a minimum Phred quality score of 20 and merged overlapping paired-end reads in Skewer. Filtered reads were aligned to the saltmarsh sparrow reference genome using *bwa-*mem 0.7.4 (Li and Durbin 2009) with default settings. Alignment statistics were obtained using qualimap version 2.1.1 (Okonechnikov et al. 2015). The average alignment rate across all individuals was 99.5 ± 0.20. After aligning sequences to the saltmarsh sparrow reference genome, depth of coverage ranged from 4.9 to 34X.

BAM files were sorted and indexed using Samtools version 1.3 (Li et al. 2009) and PCR duplicates were marked with Picard Tools version 2.1.1 (http://picard.sourceforge.net). We realigned around indels and fixed mate-pairs using GATK version 3.5 (McKenna et al. 2010). SNP variant discovery and genotyping for the 20 re-sequenced individuals was performed using the unified genotyper module in GATK. We used the following filtering parameters to remove variants: QD<2, FS>40.0, MQ<20.0 and HaplotypeScore>12.0. Variants that were not bi-allelic, had minor allele frequencies less than 5%, mean coverage less than 3X or with coverage greater than two standard deviations above the mean, and more than 20% missing data across all individuals were additionally filtered from the data set. This resulted in 7,240,443 SNPs across the genome. Using mapping results from Satsuma, we removed all SNPs located on the Z chromosome to avoid any bias that may be introduced by analyzing a mix of male and female individuals, resulting in a final data set containing 6,256,980 SNPs.

### Population Genomics

Principle component analysis (PCA) was performed on all SNPs using the SNPRELATE package in R (R core Team, 2016). We identified divergent regions of the genome by calculating F*_ST_* values for non-overlapping 100 kb using VCFtools version 0.1.14 (Danecek et al. 2011). Descriptive statistics for the 100 kb windows, including the number of fixed SNPs, the proportion of fixed SNPs in coding regions, nucleotide diversity (*π*), Tajima’s D, mean observed heterozygosity (*H_obs_*), and mean expected heterozygosity (*H_exp_*) were calculated using VCFtools and R. We calculated *D_xy_* using a custom python script (S. Martin, https://github.com/simonhmartin/genomics_general). Divergent peaks were visualized using Manhattan plots, which were constructed using the R package qqman. We discarded regions with less than two windows and windows with less than 10 SNPs.

### Characterizing Divergence between Nelson’s and Saltmarsh Sparrows

To fully assess genome-wide patterns of differentiation between saltmarsh and Nelson’s sparrows and to identify potential genes that play a role in adaptive divergence, we employed a multi-step approach to identifying and characterizing candidate regions of interest. First, we defined candidate genomic regions under selection if they contained window-based F*_ST_* estimates above the 99^th^ percentile of the empirical distribution. Second, we estimated the density of fixed SNPs (*d_f_*) in the same 100 kb windows following Ellegren et al. (2012) and identified windows above the 99^th^ percentile of the d*_f_* distribution. For both approaches, elevated windows were inspected in Genious version 9.1.5 (Kearse et al. 2012). We compiled a list of gene models within 50 kb of each elevated region and obtained information on these annotations from the UniProt database (http://www.uniprot.org/). We preformed GO analyses of divergent windows (Table 1) using the Web-based GOfinch tool (http://bioinformatics.iah.ac.uk/tools/Gofinch). Lastly, we compiled a list of candidate genes hypothesized to be important for tidal marsh adaptations, including genes linked to reproductive timing (circadian rhythm genes), osmotic regulation, salt marsh melanism, and bill morphology. Genes were chosen based on previous research done in this system (Walsh et al. 2016, 2017) and a literature review. We estimated F*_ST_* and d*_f_* for each candidate gene of interest plus 20 kb upstream and downstream of the gene, and compared our divergence estimates to the genome-wide average.

**Table 1:** Candidate genes linked to tidal marsh adaptations. Candidate regions are housed under windows exhibiting elevated divergence, assessed as regions with both F*_ST_* estimates and d*_f_* estimates higher than the 99^th^ percentile of the empirical distribution. Table includes information on location (scaffold and window start position), the number of SNPs contained within each window, mean F*_ST_,* the number of fixed SNPs, and gene name, and putative adaptive function. The full list of genes housed in elevated gene regions is found in Table S7 of Supporting Information.

| Scaffold | Window Start Position | Number of Variants | Mean Fst | Number of Fixed SNPs | Candidate Gene | Putative Adaptive Function |
| --- | --- | --- | --- | --- | --- | --- |
| scaffold_10 | 1,300,001 | 657 | 0.602402 | 223 | MSMB | Specific receptors for this protein are found on spermatozoa |
| scaffold_10 | 1,300,001 | 657 | 0.602402 | 223 | NPY4R | Blood circulation, chemical synaptic transmission, digestion, feeding behavior, neuropeptide signaling pathway |
| scaffold_10 | 1,300,001 | 657 | 0.602402 | 223 | WASHC2C | negative regulation of barbed-end actin filament capping, protein transport, regulation of substrate adhesion-dependent cell spreading, retrograde transport |
| scaffold_11 | 600,001 | 316 | 0.585958 | 144 | ATP1B1 | Catalyzes the hydrolysis of ATP coupled with the exchange of Na<sup>+</sup> and K<sup>+</sup> ions across the plasma membrane. |
| scaffold_11 | 700,001 | 288 | 0.66892 | 144 | SLC19A2 | High-affinity transporter for the intake of thiamine |
| scaffold_115 | 500,001 | 226 | 0.742673 | 147 | CAMK2D | Cellular response to calcium ion, MAPK cascade, negative regulation of sodium ion transmembrane transport, nervous system development, regulation of cellular response to heat, calmodulin-dependent protein kinase activity, sodium channel inhibitor activity |
| scaffold_115 | 500,001 | 226 | 0.742673 | 147 | NPFFR2 | Cellular response to hormone stimulus, regulation of cAMP-dependent protein kinase activity, regulation of MAPK cascade |
| scaffold_115 | 500,001 | 226 | 0.742673 | 147 | OVGP1 | Negative regulation of binding sperm to zona pellucida, single fertilization |
| scaffold_115 | 700,001 | 235 | 0.628877 | 127 | ALB | Sodium-independent organic ion transport, retina homeostasis, bile acid and bile salt transport |
| scaffold_115 | 700,001 | 235 | 0.628877 | 127 | ANKRD17 | Blood vessel maturation, defense response to bacterium, innate immune response, negative regulation of smooth muscle cell differentiation, regulation of DNA replication |
| scaffold_115 | 900,001 | 243 | 0.673404 | 130 | CXCL5 | Immune response, inflammatory response |
| scaffold_115 | 900,001 | 243 | 0.673404 | 130 | CCNI | Spermatogenesis and regulation of cell cycle |
| scaffold_115 | 1,800,001 | 418 | 0.594823 | 144 | PCDH18 | Brain development, cell adhesion, homophilic cell adhesion, nervous system development |
| scaffold_215 | 100,001 | 238 | 0.693515 | 142 | ANO2 | Calcium-activated chloride channel (CaCC) which may play a role in olfactory signal transduction |
| scaffold_261 | 1 | 368 | 0.534709 | 141 | KRTCAP 3 | Keratinocyte-associated protein 3, integral component of membrane |
| scaffold_35 | 4,500,001 | 830 | 0.530141 | 203 | KCNS1 | Potassium ion transport |
| scaffold_35 | 4,500,001 | 830 | 0.530141 | 203 | DTNBP1 | Regulates dopamine receptor signaling pathway, blood coagulation, melanosome organization and neuronal development. |
| scaffold_44 | 5,700,001 | 209 | 0.721474 | 131 | CRY1 | Transcriptional repressor which forms a core component of the circadian clock |
| scaffold_44 | 5,700,001 | 209 | 0.721474 | 131 | TMEM26 3 | Transmembrane protein |
| scaffold_44 | 6,400,001 | 229 | 0.766646 | 156 | SLC41A2 | Acts as a plasma-membrane magnesium transporter |

## Results

The final reference genome assembly generated by ALLPATHS-LG consisted of 44080 contigs with an N50 of 66.4 kb and 2672 scaffolds with an N50 of 8.427 Mb. Contig length was 1.03 Gb and the total scaffold length was 1.07 Gb. Based on a 1.0 Gb genome, sequencing coverage for the assembly was 83X. Statistics for the final assembly are included in supplementary table S2. We assessed the completeness of our reference assembly by searching for a vertebrate set of 3023 single copy orthologs using BUSCO version 1.2 (Simão et al. 2015). Our final saltmarsh sparrow reference genome contained a single and complete copy of 83.6% of the genes in the vertebrate set and a partial copy of an additional 7.5% of the genes in this set. We found 0.5% of the BUSCO vertebrate genes more than once within the reference genome and 8.8% of the BUSCO genes were missing from the saltmarsh sparrow reference.

Populations of saltmarsh and Nelson’s sparrows exhibited high levels of genome-wide divergence. Individual sparrows clustered strongly by species in a PCA based on 7.2 million SNPs, with PC axis one explaining 40.9% of the observed variation (Supporting Information; Figure S1). Genome-wide average *F_ST_* (across all autosomal SNPs) was 0.32 (Figure 2a; Supporting Information; Figure S2), indicating high baseline divergence between these two species. Genome-wide estimates of D*_xy_* showed a similar pattern to *F_ST_* (Figure 2b). We identified 234,508 fixed SNPs between saltmarsh and Nelson’s sparrows, which appear to be uniformly distributed across the genome (1.26% of which were in coding regions; Figure 2c).

**Figure 2:**
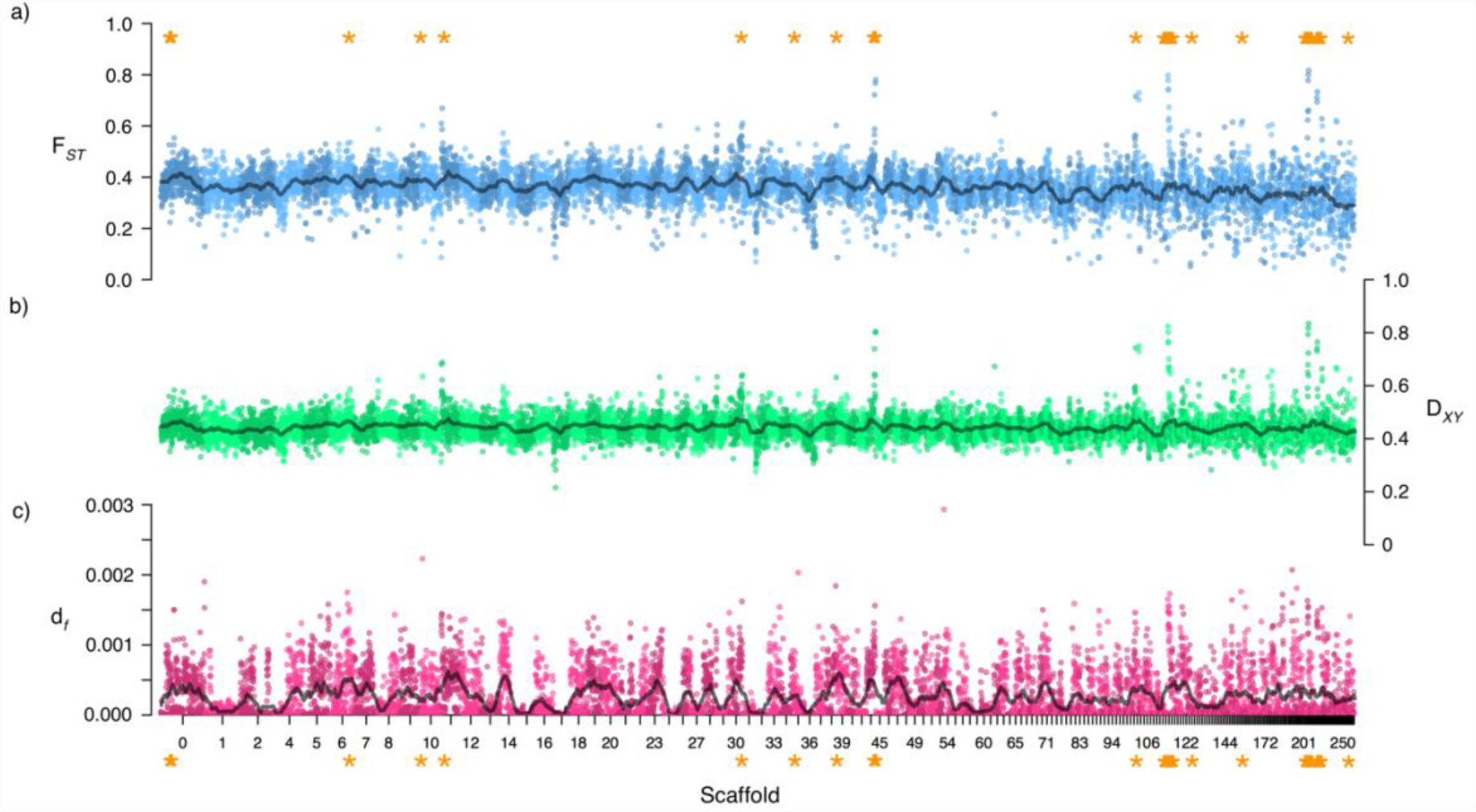
Genome-wide patterns of divergence between Saltmarsh and Nelson’s Sparrows. Panel a: genome-wide estimates of F*_ST_*; Panel b: genome-wide estimates of Dxy; Panel c: density of fixed SNPs across the genome. All results are averaged over 100 kb windows and are presented for the largest scaffolds. Elevated windows (regions of the genome with both *F_ST_* estimates and *d_f_* greater than the 99th percentile of the empirical distribution) are marked with orange asterisks.

Based on our different approaches, we identified numerous regions of the genome that exhibited elevated levels of divergence (measured as either *F_ST_* or the number of fixed SNPs in a window). We identified 90 windows across 39 scaffolds with *F_ST_* estimates greater than the 99th percentile of the empirical distribution (Supporting Information; Table S3-S4). We also identified 94 windows across 42 scaffolds exhibiting values of *d_f_* greater than the 99th percentile of the overall distribution (Supporting Information; Table S5-S6). When combining these two criteria, we found 33 windows across 16 scaffolds that showed elevated divergence using both *F_ST_* estimates and the density of fixed SNPs (hereafter *elevated regions*, Supporting Information; Table S7). When comparing elevated regions to the rest of the genome, we saw higher estimates of *F_ST_*, D*_xy_* and *d_f_* inside of the elevated regions (*F_ST_* = 0.616; D*_xy_* = 0.6795; *d_f_* = 0.0013) versus outside of the elevated regions (*F_ST_* = 0.309; D*_xy_* = 0.4412; *d_f_* = 0.00022; Table 2). Within these elevated regions, we identified 19 genes with putative adaptive roles putative roles in adaptive differences between the species (Table 1, Figure 3). Of these genes, nine were linked to osmoregulatory function, two were linked to reproductive differences between the species, two were linked to immune response, and one was linked to circadian rhythm. Enrichment analyses of these genes identified several pathways, including some linked to *a priori* hypotheses: intracellular calcium activated chloride channel activity (potentially linked to osmoregulatory function; *P* = 0.0067), regulation of circadian rhythm (potentially linked to nest initiation in relation to tidal cycles; *P* = 0.023), and sodium:potassium-exchanging ATPase activity (potential link to osmoregulatory function; *P* = 0.023). Additional genes with potential tidal marsh or other adaptive functions were identified by either the *F_ST_* or *d_f_* criteria alone (see Supporting Information. Tables S3-S6). Nucleotide diversity was decreased within these elevated windows, although not statistically significantly given the high genome-wide variation (Table 2). This could be evidence for selective sweeps in these regions. Lastly, based on a literature review and a priori predictions, we identified several *a priori* candidate genes linked to putative tidal marsh adaptations (Table 3). Of these candidates, only two had an *F_ST_* value greater than the upper bound of the 95% confidence interval for genome-wide *F_ST_* (Table 3). These candidates included CRY1 (regulation of circadian rhythm: *F_ST_* = 0.676, 86 fixed SNPs) and TYRP1 (melanin biosynthetic process: F*_ST_* = 0.553, 44 fixed SNPs).

**Figure 3:**
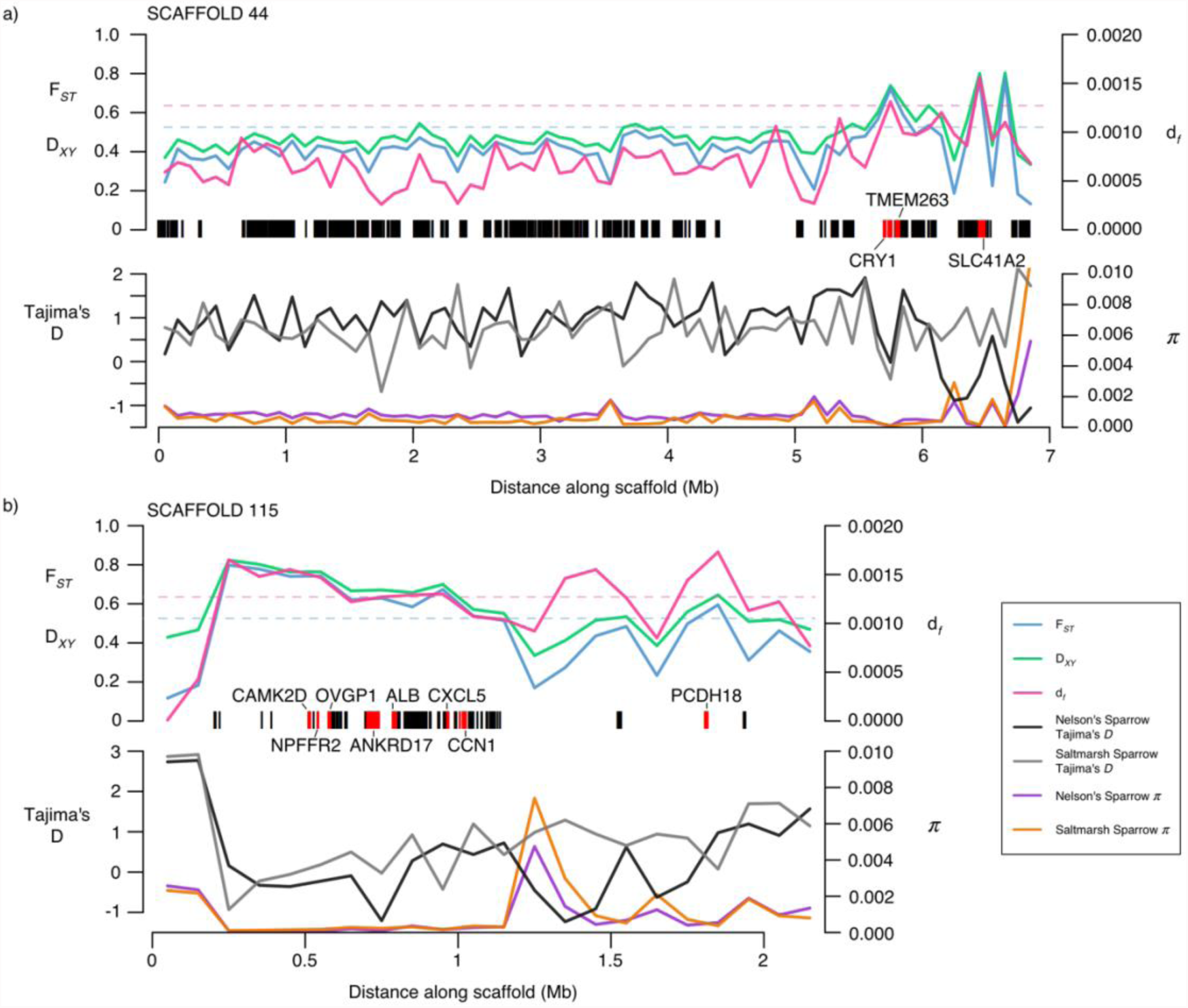
Windowed summary statistics along two scaffolds with regions of elevated divergence between Saltmarsh and Nelson’s Sparrows. Divergence is summarized by F*_ST_*, D*_XY_* and d*_f_* in 100 kb windows in the upper panel for a) scaffold 44 and b) scaffold 115. The 99% thresholds for F*_ST_* and d*_f_* are show by the blue and pink dotted lines, respectively. All annotated genes are shown by the black bars. Genes that fell within a region of elevated divergence and that we hypothesized to have functional significance within these species are highlighted in red. Nucleotide diversity (*π*) and Tajima’s D are shown for each species and scaffold along the bottom of each plot.

**Table 2:** Summary statistics for regions of the genome inside versus outside windows of elevated divergence. The mean (standard deviation) for each summary statistic is presented for the 37 windows of 100kb in length which were above the upper 99^th^ percentile of the F*_ST_* and d*_f_* distribution, along with the mean values for the rest of the autosomal genome.

|  | Inside windows |  | Outside windows |  |
| --- | --- | --- | --- | --- |
|  | Saltmarsh | Nelson's | Saltmarsh | Nelson's |
| Dxy | 0.6795 (0.0805) |  | 0.4412 (0.0428) |  |
| pi | 0.00029 (0.00017) | 0.00044 (0.00043) | 0.00101 (0.00162) | 0.00130 (0.00153) |
| Tajima's D | 0.34 (0.57) | 0.36 (0.79) | 0.33 (0.77) | 0.81 (0.61) |
| Df | 0.0013 (0.0002) |  | 0.00022 (0.00033) |  |
| $F_{ST}$ | 0.616 (0.087) | | 0.309 (0.070) | |

**Table 3:**
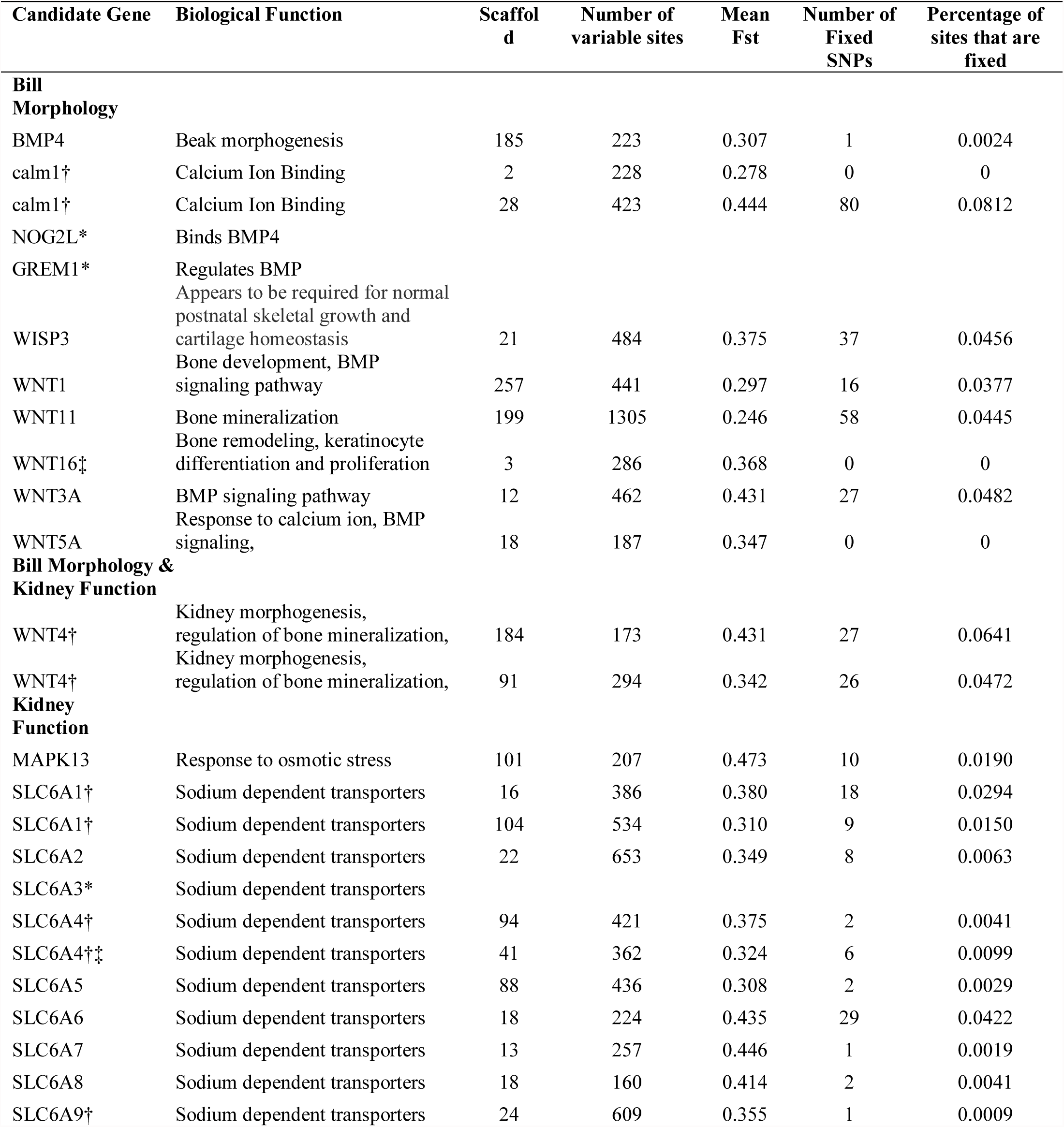
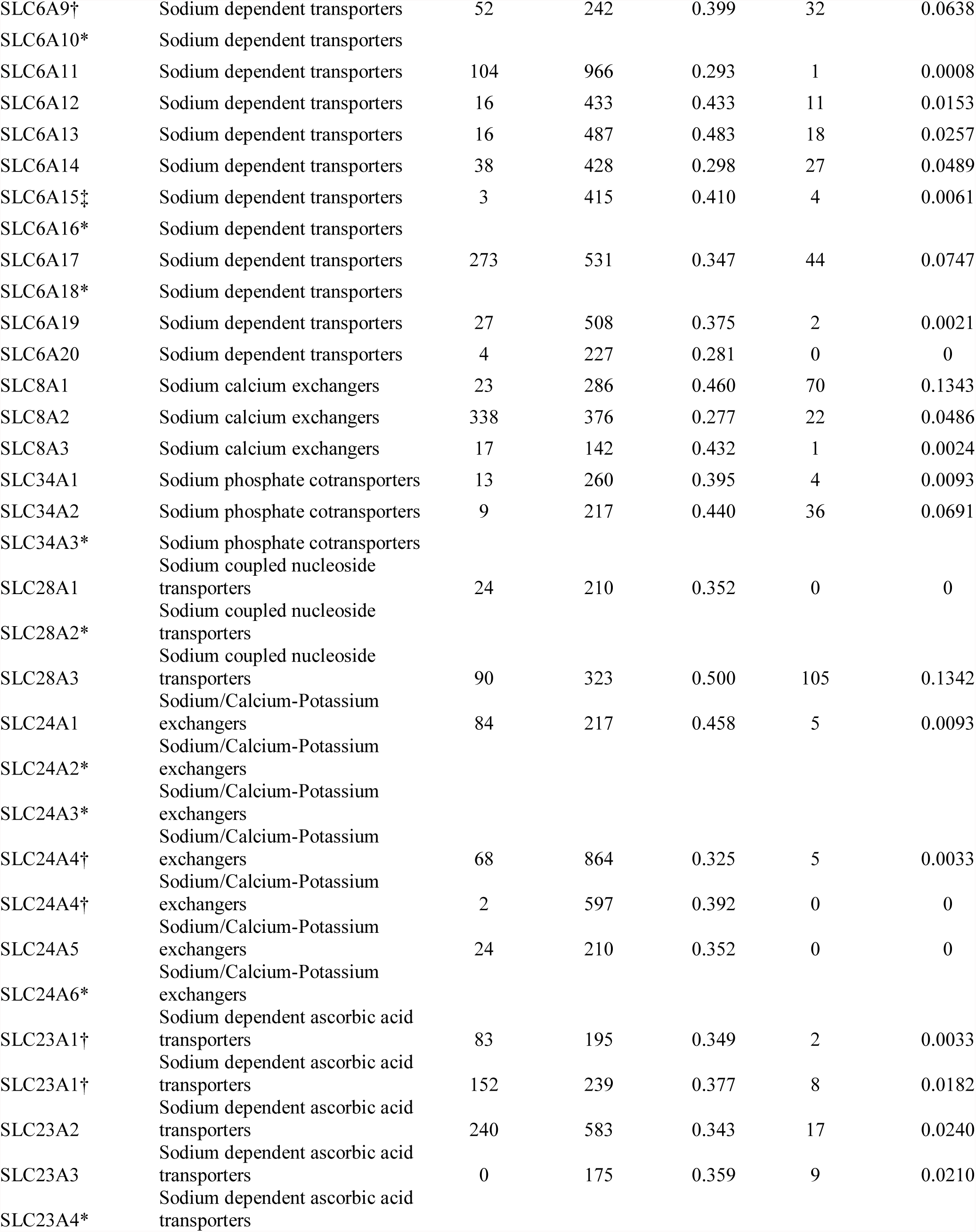
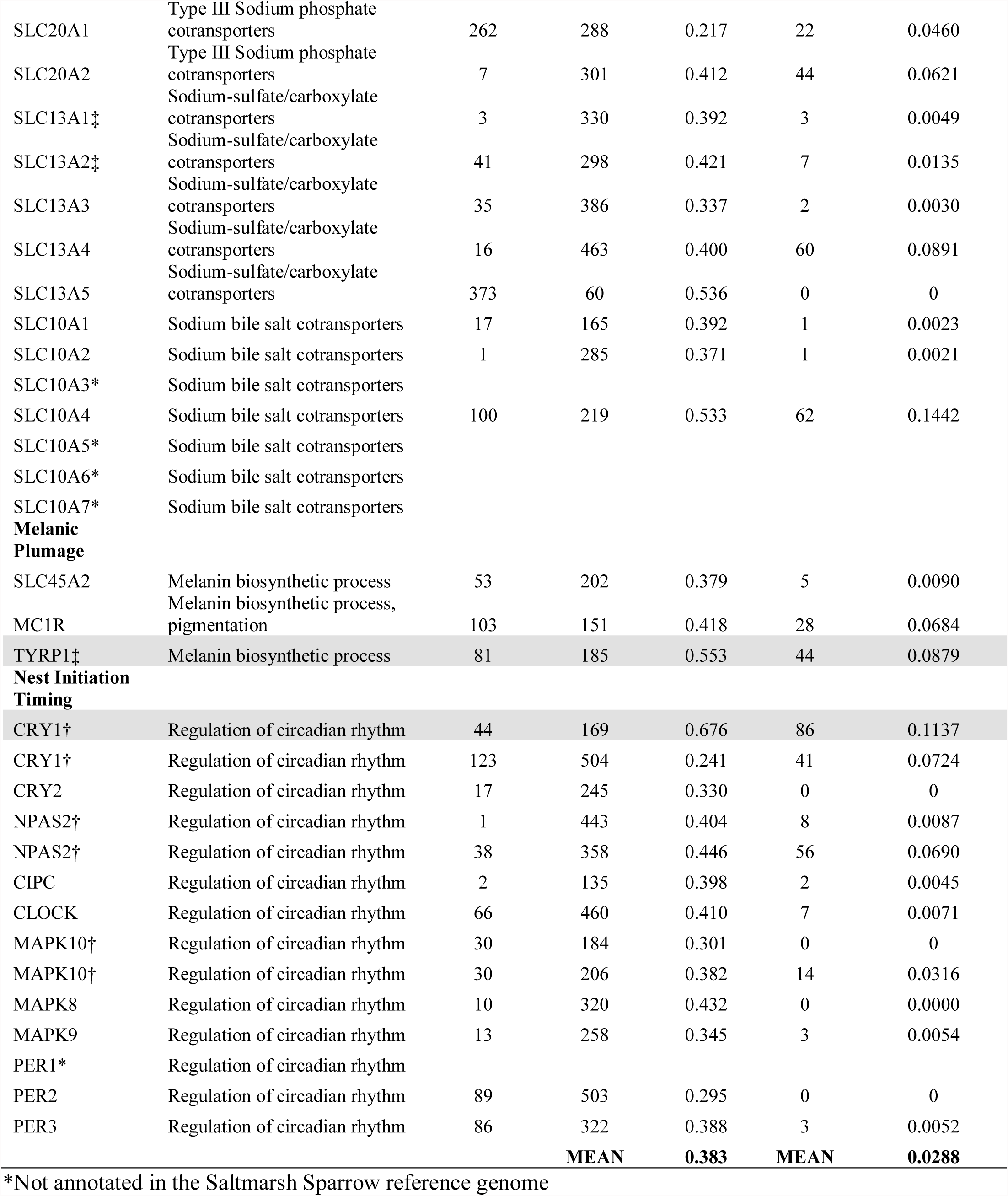
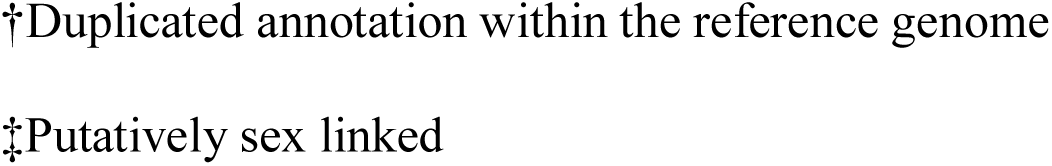
Genetic divergence measured within candidate gene regions (identified a priori). The table includes the mean F*_ST_* and number of fixed SNPs found between Nelson’s and Saltmarsh sparrows within the bounds of the coding region of each gene, plus 20 kb upstream and downstream. The percentage of sites that are fixed is calculated as the percentage of all sites in the region. Genes highlighted in grey have F*_ST_* greater than the upper bound of the 95% confidence interval.

## Discussion

Whole-genome comparisons of saltmarsh and Nelson’s sparrows revealed high baseline divergence between species (genome-wide *F_ST_* = 0.32). We identified a high density of fixed SNPs (~234,000), which appear to be uniformly distributed across the genome. These patterns share commonalities with that observed in the collared (*Ficedula albicollis*) and pied (*F. hypoleuca*) flycatcher, which exhibit high baseline divergence in allopatry (*F_ST_* = 0.357; Ellegren et al. 2012), a large number of fixed SNPs (239,745), low rates of heterospecific pairings, and fitness reductions in hybrids (Svedin et al. 2008). Conversely, flycatchers exhibit a deeper divergence time (< 2 million years; Ellegren et al. 2012) than the roughly 600 thousand years estimated between saltmarsh and Nelson’s sparrows (Rising and Avise 1993). Our findings also differ from genomic comparisons between other shallowly diverged, hybridizing taxa. For instance, Poelstra et al. (2014) and Toews et al. (2016) found only a small number of elevated regions and a small number of fixed SNPs across the entire genome (82 and 74, between carrion and hooded crows – *Corvus corone and C. cornix* – and golden-winged and blue-winged warblers – *Vermivora chrysoptera and V. cyanoptera* – respectively). Taken together, the high levels of divergence observed in these congeneric *Ammodramus* sister species may shed light on how both selective and demographic processes shape patterns of genetic differentiation.

Demographic processes, including a complex history of splitting, colonization events, and secondary contact coupled with limited contemporary gene flow are likely important factors in shaping the high baseline divergence we observed between saltmarsh and Nelson’s sparrows. It is hypothesized that the ancestral population was split by Pleistocene glaciation, resulting in an isolated interior population (Nelson’s sparrows) and a coastal refugia population (saltmarsh sparrows). Following differentiation, Nelson’s sparrow populations then spread eastward making secondary contact with coastal saltmarsh sparrow populations (Greenlaw 1993). Given repeating patterns of glaciation and retreat, it is possible that populations expanded and retracted into refugia on more than one occasion. Thus, a history of reduction in population sizes (via splitting or founder events when re-colonizing tidal marsh habitats) suggests a prominent role for drift in shaping genomic divergence between these taxa.

In addition to historical processes, patterns of contemporary gene flow may also shape the genomic landscape. Despite a 200-km hybrid zone between saltmarsh and Nelson’s sparrows (Hodgman et al. 2002; Shriver et al. 2005), our sampled individuals are predominantly from allopatric populations separated by both geographic distance and a habitat gradient that may pose a selective barrier in this system (Greenlaw et al. 1993; Walsh et al. 2015). Divergence via drift due to historical demographic processes therefore may have progressed unfettered from potential homogenizing effects of gene flow, resulting in the observed patterns of high divergence across the genomic landscape (sensu Feder et al. 2012). Support for this idea comes from a study comparing whole genomes of saltmarsh and Nelson’s sparrows from allopatric and sympatric populations (Walsh et al., in review). That study found that only about 5% of the fixed differences found in allopatry are present in sympatric populations, suggesting that when populations co-occur contemporary gene flow homogenizes all but a small portion of the genomic landscape, which likely comprises important barrier loci (loci important in reproductive isolation; Feder et al. 2012; Nosil and Feder 2012). Thus, drift – related to both historical and contemporary processes – seems to be an important factor in shaping the levels of divergence we have observed in these *Ammodramus* sparrows.

Despite a relatively high baseline of genome-wide divergence between saltmarsh and Nelson’s sparrows, we detected putative candidate regions of adaptation housed within elevated regions of differentiation (expressed both as elevated estimates of F*_ST_* and as concentrated densities of fixed SNPs). This suggests that demographic processes alone are not responsible for the observed genomic landscape and supports a role for divergent selection in shaping the observed patterns of genome-wide divergence between these taxa. Agreement on whether ecological divergence should be manifest in localized versus genome-wide differentiation is generally lacking, and it is likely that different processes are operating at different stages of the speciation continuum (Feder et al. 2012; Via 2012; Hemmer-Hansen et al. 2013). Genomic islands of divergence – discrete regions of the genome with elevated divergence harboring clusters of loci underlying ecological adaptations – have been found in ecotypes or sister species (e.g., Nosil et al. 2009, Jones et al. 2012, Hohenlohe et al. 2012, Larson et al. 2017). These islands are commonly thought to arise through divergence hitchhiking, whereby strong selection in conjunction with reduced recombination (due to linkage disequilibrium) results in coordinated evolution of multiple genes in the same genome region (Via 2012), although other mechanisms, including regions of low genetic diversity, may also explain their presence (Cruikshank and Hahn 2014). The prevalence, functional significance and underlying mechanisms of these islands of divergence are not yet fully understood (Cruikshank and Hahn 2014, Larson et al. 2017), and adaptive loci have also been found to be distributed more evenly across the genome (Strasburg et al. 2012). The pattern of high background differentiation coupled with peaks of elevated divergence distributed throughout the genomes of saltmarsh and Nelson’s sparrows is consistent with expectations for populations that diverged in allopatry, in contrast to the clustered patterns of divergence found in species that diverged with ongoing gene flow (reviewed in Harrison and Larson 2016). Under a scenario of ecological speciation, the differentiated genes within these elevated genomic regions underlie adaptations to the differential selective pressures faced by saltmarsh and Nelson’s sparrows in allopatry. We hypothesize that several of the observed genetic differences represent ecologically favored alleles in either saline or freshwater habitats, alleles underlying adaptive behaviors to tidal vs. non-tidal environments, or alleles that represent other ecological differences between the species.

Using a multi-step approach, we identified strong candidate regions for tidal marsh adaptations between saltmarsh and Nelson’s sparrows. We identified several candidates with elevated *F_ST_* estimates or numbers of fixed differences, with our most compelling candidates exhibiting elevated divergence at both of these metrics. Candidates with putative adaptive roles in tidal marshes include genes linked to osmoregulatory function, circadian rhythm, and melanin pigmentation. The genes SLC41A2 and ALB, for example, exhibited elevated *F_ST_* estimates and a high density of fixed SNPs, and have roles in solute transport pathways that may occur in osmoregulatory function. SLC41A2 is a membrane Mg^2+^ transporter. Mg^2+^ is the second most abundant cation in seawater, with SLC41A2 identified as Na^+^/Mg^2+^ exchanger that is highly expressed in the kidney of saltwater acclimated puffer fish (*Takifuga rubripes)* compared to closely related freshwater species (*T. obscurus;* Islam et al. 2013). SLC41A2 has also been previously found to exhibit reduced introgression and increased selection across the saltmarsh Nelson’s sparrow hybrid zone (Walsh et al. 2016), offering further support for an adaptive role for this gene. ALB is linked to the regulation of osmotic pressure of blood plasma and is a gene that was found to be under positive selection in the saline tolerant crab-eating frog (*Fejervarya cancrivora*) compared to a morphologically similar, saline intolerant sister species (Shao et al. 2015).

Another candidate gene linked to tidal marsh adaptation identified in this study was CRY1, which is an important component of the circadian clock. In salt marshes, flooding affects nests during the highest spring tides; during this time the entire marsh is flooded and nests can be inundated with water for an hour or two (Gjerdrum et al. 2008). Monthly tidal flooding, therefore, is the leading cause of nest failure and an important driver of overall population trajectories for tidal marsh sparrows (Greenlaw and Rising, 1994; Shriver et al. 2007); synchronizing the 24-26-day nesting cycle with the approximate 28-day monthly tidal cycle is critical for individual fitness. Saltmarsh sparrows have greater nesting synchrony with tidal cycles compared to Nelson’s sparrows and this synchrony is associated with increased nesting success (Shriver et al. 2007; Walsh et al. 2016). Biological clocks are important for ensuring that the passage of time is synchronized with periodic environmental events (Kumar et al. 2010); as such, it is reasonable to hypothesize that CRY1 may play a role in the divergent nesting synchrony observed between these species. Expression patterns of CRY1 were shown to fluctuate with lunar periodicity in a lunar-synchronized spawner, the goldlined spinefoot (*Siganus guttatus*; Ikegami et al. 2014), supporting a link between CRY1 and lunar cycles.

The final putative candidate for tidal marsh adaptation that we identified was TYRP1, which is an important component of the melanin biosynthesis pathway and has been found to play an important role in determining plumage color in birds (Xu et al. 2013; Minvielle et al. 2009; Bourgeois et al. 2016). This finding supports previous work identifying SLC45A2 (another gene associated with plumage melanism) to be under divergent selection in this system (Walsh et al. 2016). Saltmarsh and Nelson’s sparrows have subtle plumage differences – with saltmarsh sparrows showing darker breast and flank streaking and face coloration than Nelson’s sparrows (Shriver et al. 2007) – consistent with phenotypic patterns observed in other vertebrates spanning tidal marsh gradients (Grinnell 1913; Luttrell et al. 2014). Tidal marsh taxa are grayer or blacker than their upland relatives, due to a greater expression of eumelanin relative to phaeomelanin (Greenberg and Droege 1990). This salt marsh melanism is thought to confer adaptive benefits to tidal marsh birds either via enhanced predator avoidance (from background matching with the gray-black marsh mud; Greenberg and Droege 1990) or resistance to feather degradation (increased melanin concentrations slow degradation rates by salt-tolerant feather bacteria; Goldstein et al. 2004).

In addition to candidate genes that were in line with our *a priori* predictions for tidal marsh adaptations, we also identified several genes under selection that are related to spermatogenesis. Differences in mating strategies in saltmarsh and Nelson’s sparrows (Greenlaw 1993) coupled with strong assortative mating in the hybrid zone (Walsh et al. in press) support a role for pre-mating barriers, which could include sperm competition or incompatibilities between the species. These candidate regions, coupled with the osmoregulatory and circadian rhythm candidates described above, will provide directions for future research in this system, including candidate gene work in more individuals across habitat types. Further, the full suite of genes identified by either *F_ST_* or *d_f_* criteria alone (Supporting Information; Tables S3-S6) provide additional putative candidates for further investigation.

The candidates for osmoregulatory response, circadian rhythm, and melanistic plumage described above are likely representative of important lineage specific adaptations and add to a body of empirical evidence describing underlying mechanisms driving adaptation to harsh environments (Wu et al. 2014; Yang et al. 2016; Tong et al. 2017). Our findings, in particular, contribute to a growing list of candidate genes linked to salt tolerance and osmoregulation (Kahle et al. 2010; Ferchaud et al. 2014; Gibbons et al. 2017), suggesting that modifications to several pathways can result in adaptations to saline environments. Lastly, our results demonstrate the effects of both ecological divergence and drift in driving high baseline levels of differentiation between two closely related sister taxa. A high proportion of fixed SNPs distributed across the genomes of saltmarsh and Nelson’s sparrows offers new perspectives into processes shaping the genomic landscape and offers empirical evidence for the role of ecological divergence and demography in shaping evolutionary processes and genetic variation between these taxa.

## Acknowledgements

Funding for this project was provided by the New Hampshire Agricultural Experiment Station through a USDA National Institute of Food and Agriculture McIntire-Stennis Project # 225575, the Maine Association of Wetland Scientists, and the Robert A. Chase’45 and Ann Parker Chase’46 Faculty Fund at the University of New Hampshire. This is Scientific Contribution Number XXXX of the New Hampshire Agricultural Experiment Station. We thank Kazufusa Okamoto for transcriptome assembly and Stephen Simpson for library development of resequenced genomes.

## Data Accessibility

The Illumina reads are available from the sequence reads archive (study number: TBD). Our SNP dataset and the saltmarsh sparrow reference genome is available from the Dryad Data Repository (doi: TBD).

## Author Contributions

A.I.K. conceived and designed this study, with input from J.W., G.V.C., and M.D.M. J.W. and A.I.K. collected samples. J.W. and G.V.C. analyzed the data. J.W., G.V.C. and A.I.K. wrote the manuscript. M.D.M. provided guidance with bioinformatics analyses. W.K.T. lead the transcriptome development and genome resequencing process. All authors read and approved the final manuscript.

